# Random attention can explain apparent object choice behavior in free-walking blowflies

**DOI:** 10.1101/2021.11.15.468665

**Authors:** José Monteagudo, Martin Egelhaaf, Jens Peter Lindemann

## Abstract

Flies are often observed to approach dark objects. To a naïve observer they seem to pay selective attention to one out of several objects although previous research identified a reflex-like fixation behavior integrating responses to all objects as possible underlying mechanism. In a combination of behavioral experiments and computational modelling, we investigate the choice behavior of flies freely walking towards an arrangement of two objects placed at a variable distance from each other. The walking trajectories are oriented towards one of the objects much earlier than predicted by a simple reactive model. We show that object choice can be explained by a continuous control scheme in combination with a mechanism randomly responding to the position of each object according to a stochastic process. Although this may be viewed as a special form of an attention-like mechanism, the model does not require an explicit decision mechanism or a memory for the drawn decision.

**Summary Statement:** Walking blowflies apparently choose one of two objects to approach. In model simulations, a fixation scheme combined with random attention replicates this behavior without an explicit decision mechanism.

## Introduction

Despite being known mostly for their flight behavior, flies often also explore their environment by walking. During such explorations, flies have frequently been observed to approach distinct dark objects (Wehner 1972). For a human observer individual walks in presence of multiple objects seem to indicate selective attention to a single target, because on individual trajectories the flies usually turn towards one of the objects in a quick turn apparently indicating a decision to approach the chosen target.

In flies, the mechanisms underlying such object-related orientation behavior have mostly been studied in tethered flight. In such experimental paradigms with a single object, such as a vertical bar, flies tend to fixate this object for most of the time in the frontal visual field. They accomplish this by generating a torque that depends on the object’s azimuthal position and is directed towards it. The strength of this turning response peaks when the object is at a fronto-lateral position (Reichardt 1973). Consequently, the object moves into the frontal visual field and is stabilized in front of the animal. A similar mechanism was proposed for walking flies (Horn and Wehner 1975, Horn 1978).

When a fly is confronted not with just one object but with two objects in a closed-loop tethered flight scenario, it fixates one of the objects unless the objects are close to each other; then the midpoint between them is fixated (Reichardt 1973, Poggio and Reichardt 1973, Reichardt and Poggio 1975). These experiments led to the conclusion that, if confronted with two objects, flies react simultaneously to both by summing the torque responses predicted to be induced by each individual object according to the fixation characteristic determined for an object when presented in isolation. This additive concept is consistent with the observation that, when the objects are separated by 60° or more (i.e. the distance between their azimuthal positions at which maximum torque is generated), one object could be fixated frontally, because the torque generated by the more lateral object is smaller in this case than the torque generated by the more frontal object. If the objects were less far apart, both generate similar torques resulting in the animal orienting itself towards the mid-line between the two objects (Reichardt 1973, Reichardt and Poggio 1976). Similarly, the behavioral data obtained from walking flies facing two or three objects could be explained by adding the responses elicited by the objects when presented individually (Horn and Wehner 1975). All these conclusions were based on the overall behavioral performance averaged over many flights or walks and animals.

However, when scrutinizing the time-dependent performance of flies during individual flights in an open-loop two-object paradigm with one bar in each half of the visual field at corresponding positions, the torque responses of the two bars did not cancel out if these were oscillated synchronously in anti-phase in contrast to predictions by the additive fixation model. Instead, the flies responded as if only one bar were present, apparently ignoring the other temporarily before switching after some time to respond to the other bar (Wolf and Heisenberg 1980). This behavior has been interpreted as a consequence of selective attention and attention switches between the objects. These analyses were performed under tethered flight conditions, where the animal could not approach the objects, essentially simulating a condition in which the objects were at an infinite distance, a situation occurring in a fly’s real life only under very special conditions.

To overcome this limitation, we performed behavioral experiments on free-walking flies in a specially designed object choice paradigm. The use of a walking paradigm made it possible to monitor object-oriented decision behavior of largely unconstrained flies using video techniques, an approach hardly possible with free-flying flies. Some work has already been done on walking flies before to address which object parameters appear to be attractive to flies resulting in frequent approaches, such as the height and width of a bar (Wehner 1972) as well as, based on studies in a virtual reality closed-loop paradigm, its distance as inferred from relative motion cues (Schuster et al. 2002). Further systems analysis suggested that flies were attracted by fast-moving bars in general (Mronz 2004). Despite this evidence on object preferences of walking flies, a comprehensive concept of how individual flies select an object for approach in a choice situation is still lacking.

To address this issue, we developed a behavioral paradigm in which object selection behavior of freely walking blowflies (*Lucilia spec.*) could be recorded for various spatial relationships between the objects and the fly. The time-dependent object selection and fixation behavior is then related to the performance of the classical additive fixation model (see above) as well as a newly developed model that combines the simultaneously summed response to individual objects with the ability to stochastically ‘ignore’ one of them. In contrast to the classical additive fixation model, this new model is sufficient to reproduce the choice behavior of walking flies as characterized by our experiments. It further reveals that apparent choice behavior can be explained without the need of an explicit decision module.

## Methods

### Animals and animal preparation

We did our experimental analysis on female blowflies (*Lucilia spec.)* bred in our laboratory. Animals were captured 1-3 days after hatching, briefly anesthetized with CO_2_, and prevented from flying by placing a drop of wax on the wing joints. The prepared animals were kept in a cage with *ad libitum* access to sugar and water.

### Experimental setup

Our experimental setup (Fig. 1) consisted of an irregularly pentagonal arena (for dimensions see Fig. 1 A), constructed of canvas frames covered with a random cloud pattern with spatial statistics characterized by a 1/f spectrum.

**Figure 1:**
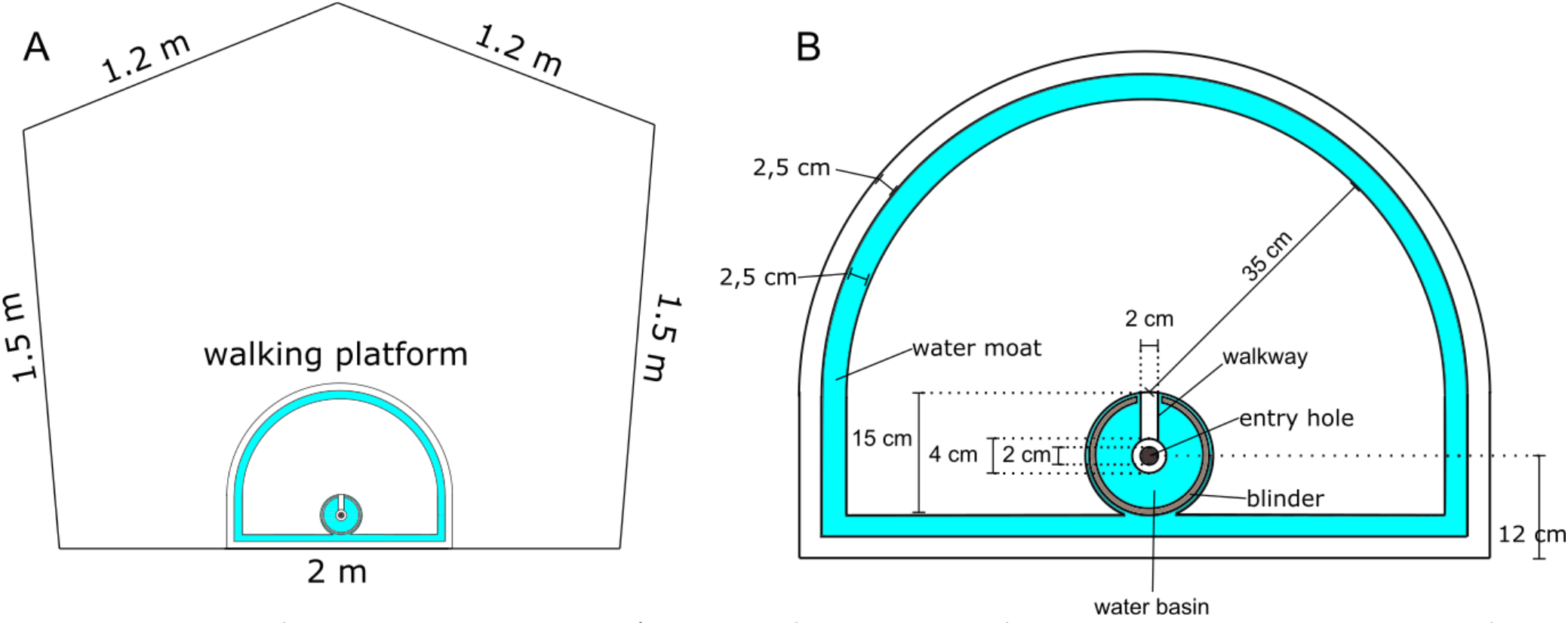
Sketch of experimental setup. A) Sketch of a top view of the arena walls and position of the walking platform. Arena walls are 1 m high C) Sketch of the walking platform. Blue color marks the position of the water moat being 0.5 cm deep. The blinder is 3 cm high and leaves 0.5 cm to both sides of the walkway.

A walking platform was placed within the arena, centered against the 2 m long wall. The walking platform was cut from white PVC (for dimensions see Fig. 1 B) with an entry hole through which animals were individually introduced into the setup. To control the initial direction of orientation of the animal relative to the experimental arena when the objects get in sight, the entrance hole is surrounded by a water basin, and the view into the test arena is initially occluded by a blinder surrounding it. Animals had to walk into the arena via a narrow walkway connecting the entry hole to the experimental area. By this arrangement we achieved a relatively uniform walking direction and body orientation of the animals when entering the arena. A water moat surrounding the walking platform keeps animals on the platform. Both the water moat and the basin are connected and are 0.5 cm deep. The walking platform was elevated by 5.5 cm from the supporting table to hide construction details of the setup from the animal’s view.

Cylinders of 8.2 cm diameter and 20 cm height were placed at a distance of 60 cm from the end of the walkway. The cylinders were placed at three different angular positions relative to the midline of the walkway as a reference line (0°). If two objects were placed in the arena, one was placed in the right half of the arena and the other one in the left half.

The arena was illuminated indirectly through a diffusion screen by 6 white LED lamps (Marathon MultiLED, GS Vitec GmbH, Gelnhausen, Germany) placed above the arena. The resulting soft light reduced shadows and allowed for automated tracking of the walking flies.

The behavior of walking blowflies was recorded at 90 fps using a camera (Basler acA 2040-90um, Basler AG, Ahrensburg, Germany) placed above the arena and a custom-made recording software based on the Pylon 4 software suite (Basler AG, Ahrensburg, Germany). Animals were tracked on the video using the open-source software ivTrace (https://opensource.cit-ec.de/projects/ivtools). The animal’s position and orientation were automatically determined by fitting an ellipse to the body and using the orientation of the ellipse’s long axis as a proxy of its gaze direction and the center of ellipse as its position. The tracking results were reviewed and obvious misdetections were manually corrected to fit the position and orientation of the animals.

### Experimental procedure

Individual blowflies were released from below through the entry hole into the experimental setup. Flies were recorded until they reached the borders of the walking platform. If an animal attempted to take off or failed to reach the outer borders of the platform because it refused to walk, it was captured and released again. Each individual was recorded under a given stimulus condition until it reached the borders of the platform 10 times. We recorded blowflies walking under seven conditions: in the absence of objects, in presence of one object at 37°, at 60°, or at 90°, and in the presence of two objects at 37° and 60°, at 37° and 90°, or at 60° and 90°. For each condition, we recorded 10 different flies. For both the one- and the two-object conditions, the object constellation was mirrored along the symmetry axis of the walking arena between experiments according to a pseudo-random sequence, to control for any potential asymmetry in the experimental arena that might have escaped our notice.

### Characterization of walking trajectories

We defined the start of apparent object fixation as the onset of the time window where at least one of the object’s edges has been kept in the frontal visual field, i.e. within ± 30° relative to the midline of the animal, for at least 100 frames (^~^ 1.11s). In the following these phases will be called “lock-on”.

### Comparison between model and experimental data

To compare model performance and corresponding experimental data, we generated 100 walks for each condition for every model variant tested. The start location of the modelled animal was at the end of the walkway because we aimed to model the behavior of the animal once it was no longer constrained by the walkway. The exact positions and orientations were varied to match the observations in the experiments: For each walk recorded in the experiments, we started the model trajectory at the position at which the animal left the walkway and with the orientation it had in that moment.

### Modelling: Additive Fixation Model (AFM)

We attempted to qualitatively reproduce the experimental data with a model that was inspired by previous work (Poggio and Reichardt 1973, Horn and Wehner 1975). This additive fixation model (AFM) achieves the fixation behavior by summing two behavioral components (Fig. 2, gray boxes):

i. An object-induced turning behavior, generating torque depending in amplitude on the azimuthal position of the object. The corresponding characteristic function is defined as

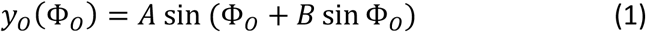

where *y*_0_ is the resulting yaw speed, and Φ_0_ the azimuth position of the object *O. A* and *B* are free parameters controlling peak yaw speed and peak position, respectively. To parameterize the characteristic curve, we fitted the average yaw speed observed for a given retinal object position using data obtained when a single object was initially present at 90<u>º</u>. The plot of yaw speed versus azimuthal position generated in this way follows an anti-symmetrical curve (cf. inset in Fig. 2). The parameters were generated by least-squares fit of Eq. (1) to the experimental data (R^2^=0.852). If two objects are present in the visual field the yaw torque resulting for each object is determined individually for its actual retinal position according to the characteristic curve. The overall object-induced yaw torque is then obtained by adding the individual torque components.
ii. A spontaneous stochastic turning tendency implemented by temporally filtered white noise fluctuations. The filter kernel was determined from the spontaneous walking behavior of the flies in our setup without objects (for details, see Monteagudo Ibaretta 2020). We determined the average fast Fourier transform (FFT) of the yaw speed for all recordings lasting at least 512 frames, allowing us to use 47 of the 100 recordings. Using the inverse fast Fourier transform (iFFT), we converted the average spectrum to a linear temporal filter kernel. Applying this filter to a white noise signal by convolution and normalizing the result to the standard deviation of the observed yaw velocities, we generated a sequence of spontaneous turns.

**Figure 2:**
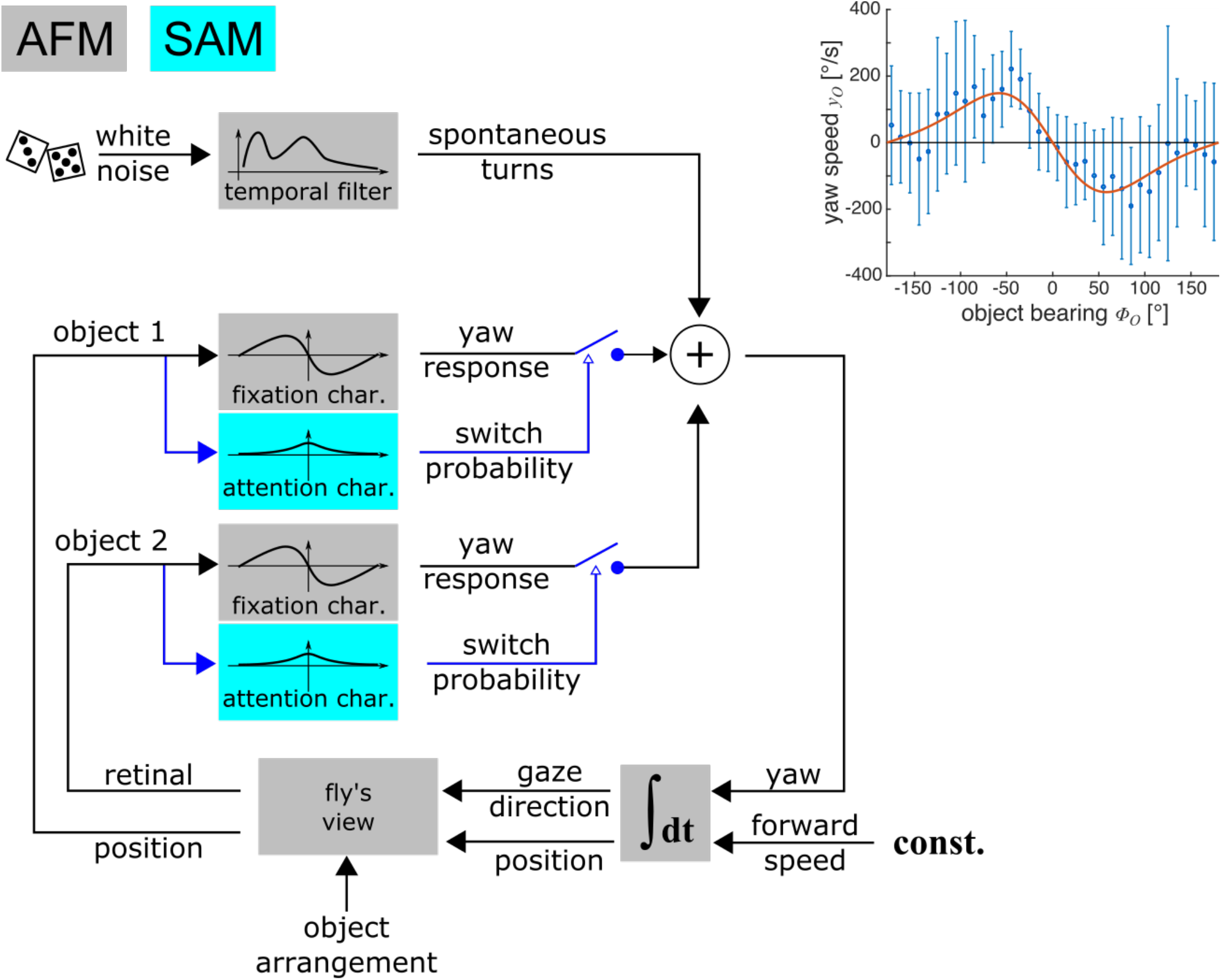
Sketch of models. Modules in gray boxes form the additive fixation model (AFM). Blue modules are extensions to add for the stochastic attention model (SAM). The AFM generates spontaneous turns during walking from temporally filtered white noise to which the torque response to both objects is added. The resulting change in yaw orientation is integrated to update the retinal positions of the objects assuming constant speed walking. In SAM the torque response to each object is added with a certain probability modulated by the attention curve. The inset shows the fixation characteristic generating the object responses as derived from experimental data: data shown in blue shows mean +− S.E.M. of the yaw speed observed when the object is at different azimuth positions, red curve shows the functional fit to the data used in the model.

We assumed a response delay of the fly of 4 frames (^~^44 ms), i.e. the object response was calculated for the object position 4 frames earlier. This delay approximates the neuronal latency observed for motion-sensitive wide-field neurons in the lobula complex of the fly (Warzecha and Egelhaaf 2000). The walking speed was set constant throughout the simulations and corresponded to the average walking speed of 6.4 cm/s of our experimental animals along their trajectories (cf. Fig. S1). The simulated trajectory was updated at a 90 Hz frequency. The orientation of the model animal was updated by stepwise integration of its yaw speed, which is in turn controlled by two components: the spontaneous turns and the object response.

### Modelling: Stochastic Attention Model (SAM)

The stochastic attention model (SAM) has the same overall structure as the AFM with just one extension, i.e. the simulated animal ignores one or both objects stochastically (Fig. 2, blue symbols). This is accomplished by switching the object responses ‘on’ or ‘off’ before contributing to the overall yaw torque following a random process modulated by the object’s azimuthal position. This random switch is applied independently in each update step and to each object in the visual field. The function describing the dependence of the switching probability on the object position will be called the attention curve and was assumed to be bell-shaped according to a von Mises distribution, i.e. the circular normal distribution. We used a modified von Mises distribution with a fixed maximum position at 0° (*μ* = 0) and scaled by systematic parameter variation:

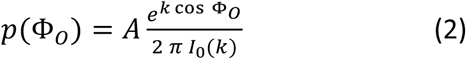

where *p*(Φ_0_) is the attention probability for the object *O* observed at the azimuthal position Φ_0_. *A* is the varied scaling factor, and *k* is the constant parameter of the zero-order modified Bessel function *I*_0_. The object response was only added if a uniformly distributed random number between 0 and 1 was below the probability according to the value of equation 2. Otherwise, the object response was set to *y*_0_ = 0 °/*s*. The attention curve generating the attention probability for a given object position was parameterized by systematic variation of the parameters *k* and *A* of eqn. (2) for 100 simulated trajectories and the similarity of the resulting model trajectories to the experimentally observed ones was assessed by visual inspection (compare Fig. 3 A and 6 A) and set *k* = 5.518 and *A* = 96.37. The resulting von Mieses curve is the circular equivalent of a Gaussian with *σ* = 25° and a maximum attention probability of *p*_*max*_ = 0.875.

**Figure 3:**
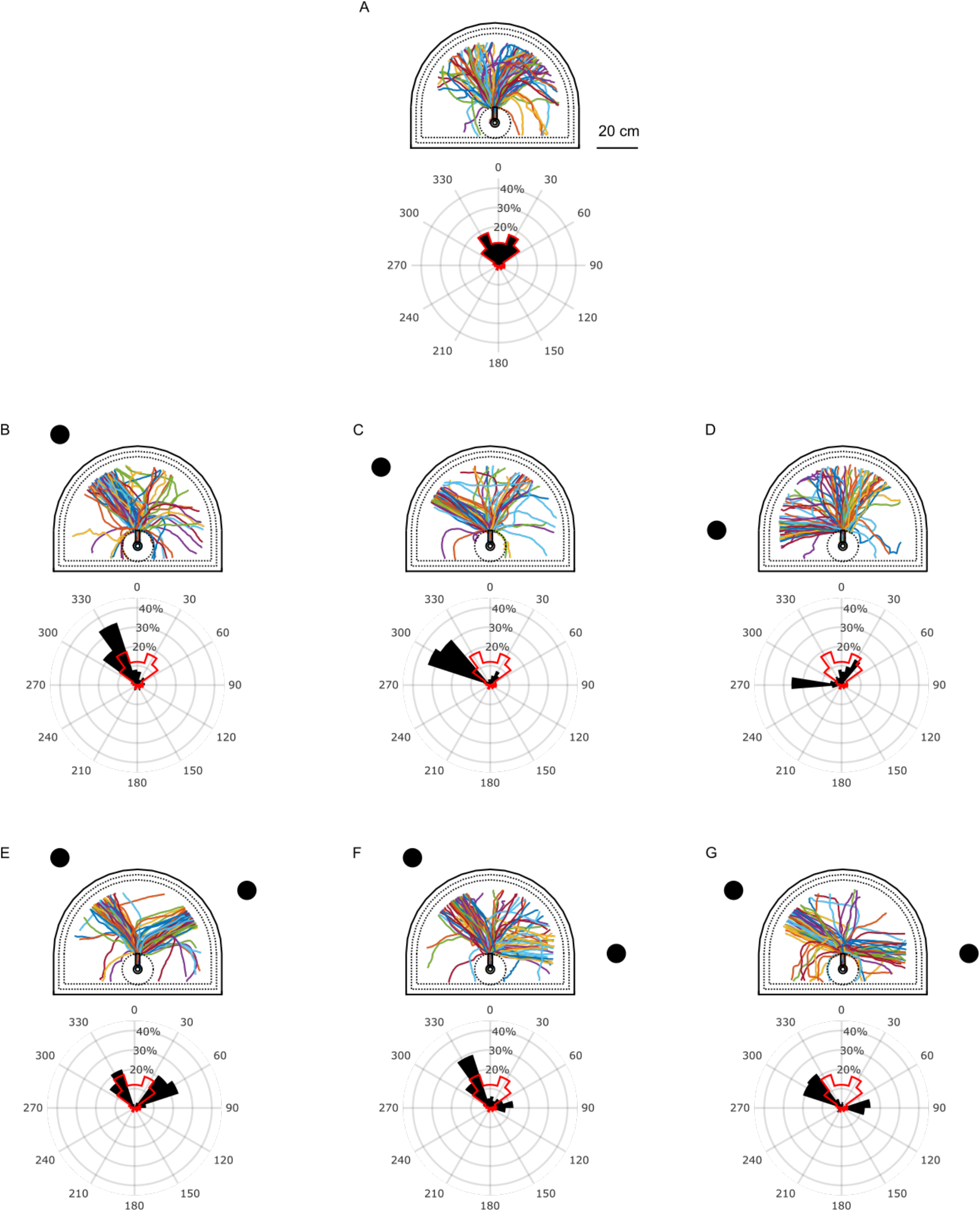
Trajectories of walking blowflies. A) In absence of any object. B-D) With one object at 37°, 60°, or 90°, and E-G) With two objects, at 37° and 60°, at 37° and 90°, or at 60° and 90° respectively. Trajectories were colored for better separability, colors do not imply any grouping. The polar histograms show the distribution of positions observed when the flies crossed a registration circle with 20 cm radius around the end of the walkway. Red silhouettes in B - G repeat the distribution of A for comparison.

**Figure 4:**
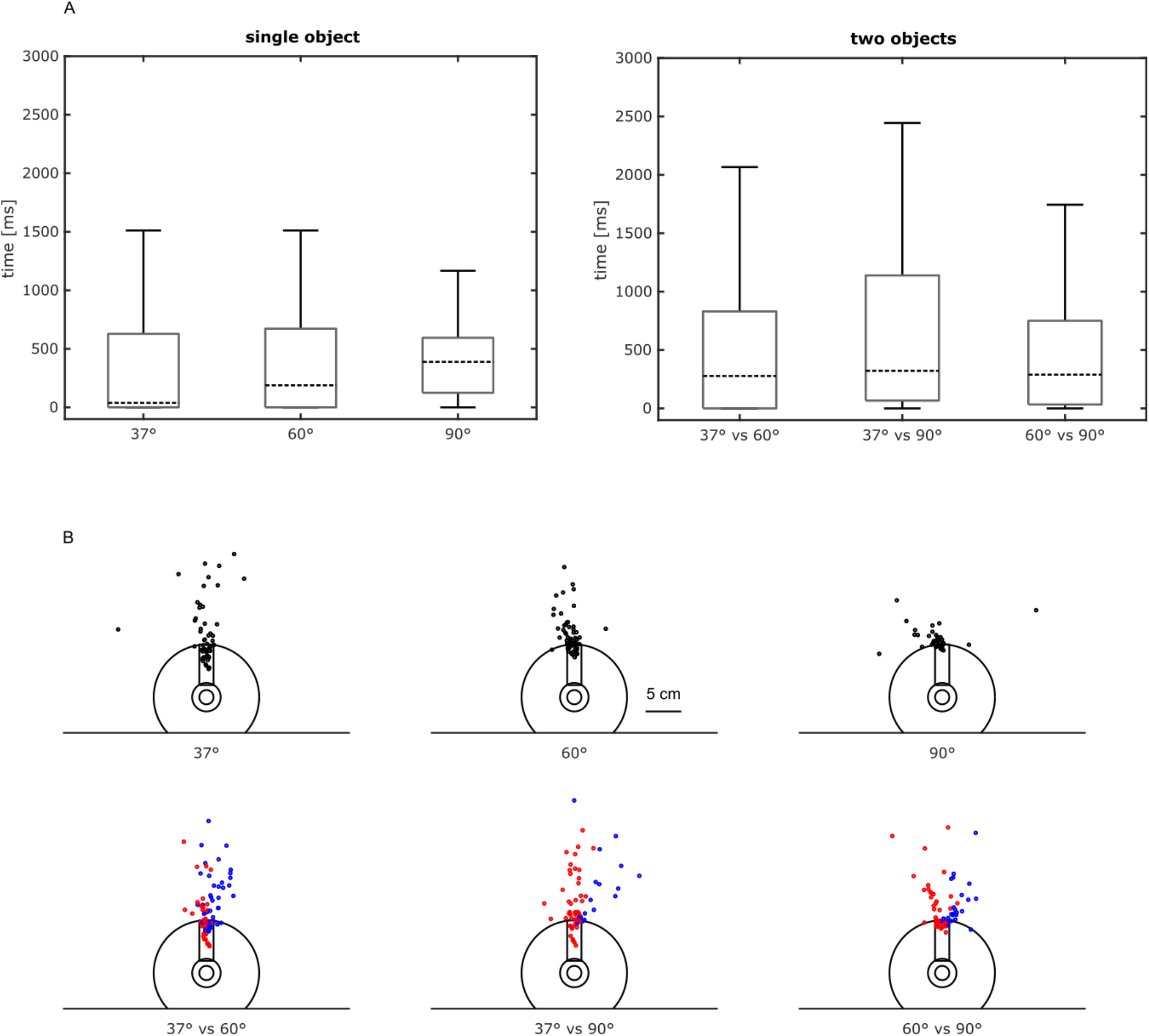
Analysis of object lock-on during walks. A) Boxplots of the time between leaving the walkway and object lock-on. A few durations exceeding the limit of the y-axis are not shown in the plot. B) Location of lock-on moment. Upper row: single objects. Lower row: two objects, red dots indicate walks approaching the object more frontal to the walkway, blue dots the one more lateral.

**Figure 5:**
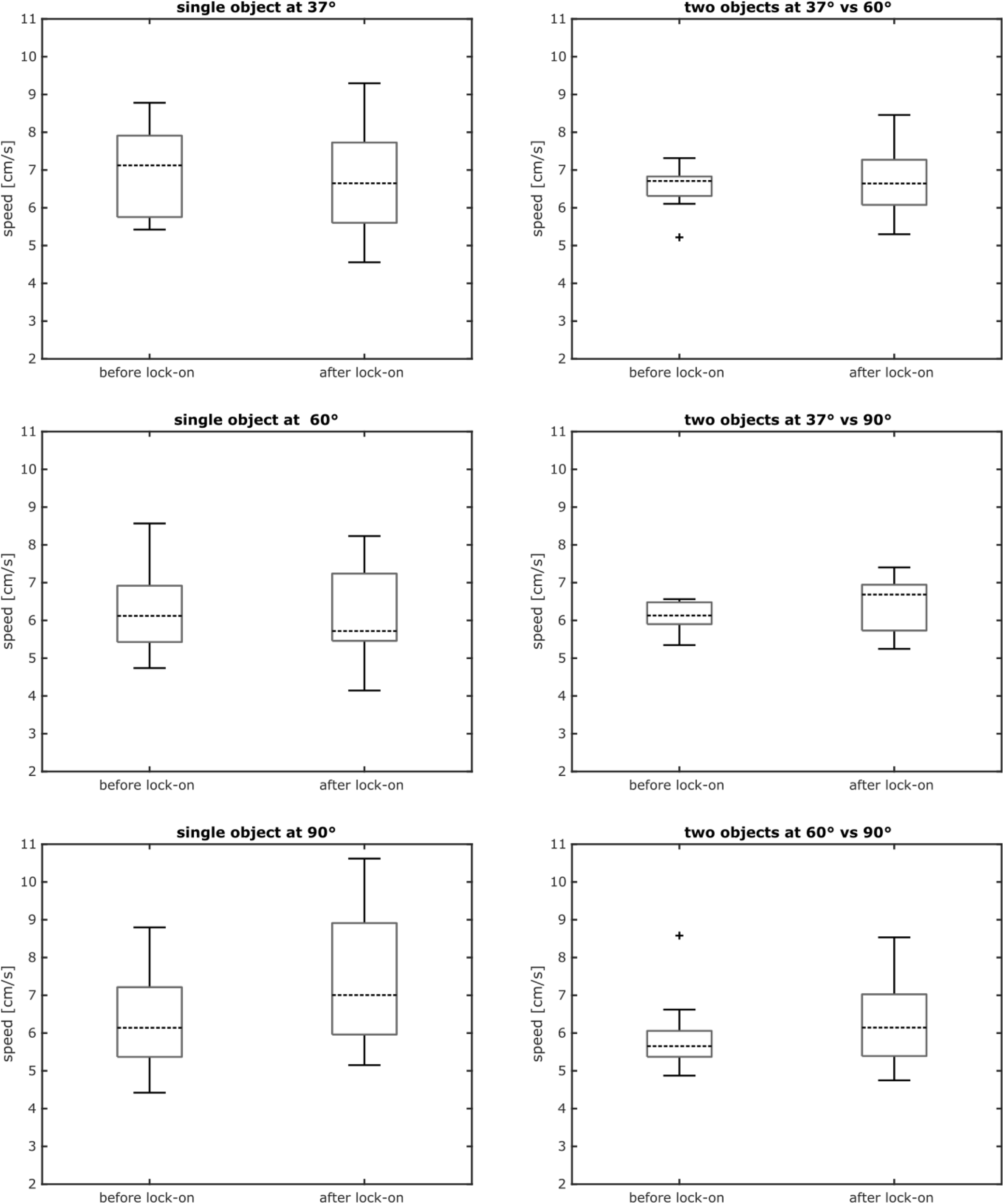
Comparison of mean walking speed of blowflies before and after lock-on to an approached object. Object constellations are given in the plot titles.

**Figure 6:**
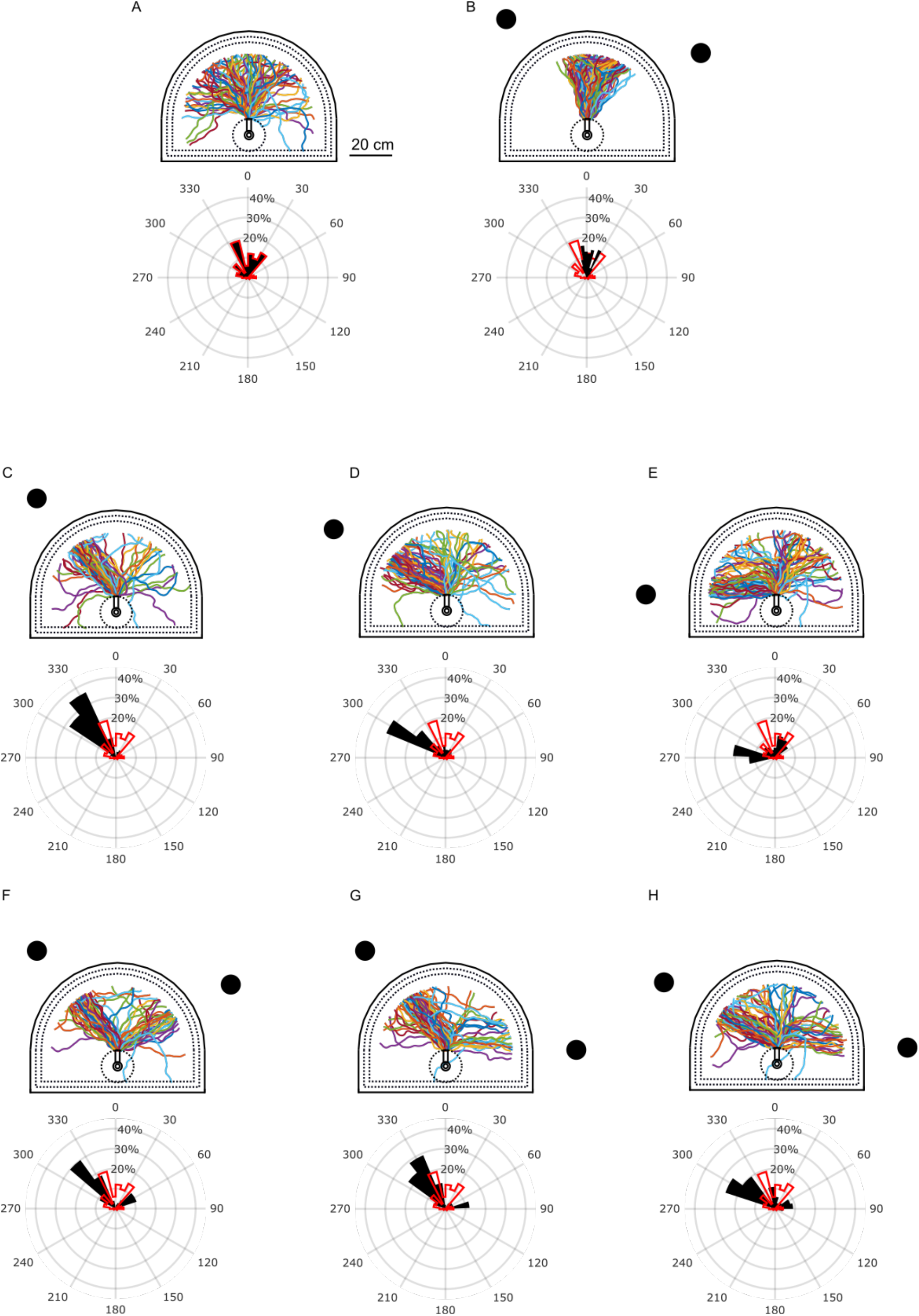
Walking trajectories generated by the model. A) Trajectories resulting from spontaneous turns in absence of any object-related response. B) Trajectories generated by the additive fixation model (AFM) without attention-like processes. C-E) Trajectories generated by the stochastic attention model (SAM) for single objects. F-H) Trajectories generated by the SAM when tested with two objects. Trajectories were colored for better separability, colors do not imply any grouping. The polar histograms show the distribution of positions observed when the model trajectories crossed a registration circle with 20 cm radius around the end of the walkway. Red silhouettes in B - H repeat the distribution of A for comparison.

## Results

### Behavioral experiments

To gain insight into the goal selection mechanisms of walking blowflies, we analyzed their free walking behavior in a two-object paradigm and developed a model reproducing key features of the observed behavior.

In an environment without any obvious visual object most walks tended to proceed in their initial walking directions along fairly straight trajectories. Only in some cases we observed flies walking on a strongly curved path or reorienting substantially between leaving the walkway and reaching the water moat (Fig. 3 A).

In the presence of a single object presented at different azimuthal positions, in most walks the animals approached the object along fairly straight paths right from the moment they enter the arena (Fig. 3 B - D). However, in some cases the animals walked in different directions on similarly straight trajectories, sometimes animals even turned away from the object after initially heading towards it. Only a few walks led to a curved trajectory.

The objects were approached with a variable probability depending on their position in the arena (see histograms in Fig. 3 B - D). An object at 37° or 60°, as seen from the initial walking direction, is approached more frequently than an object at 90°; objects at 37° and 60° are approached similarly often.

When confronted with two objects, most flies approached only one of them on a given walk, with the probability of approach of either object depending on its azimuthal position (Fig. 3 E - G). A variable proportion of flies did not approach either object or turned away after an initial object approach. Flies preferred the more frontal object over the lateral one (37° over 90°, Fig. 3 F; or 60° over 90°, Fig. 3 G), like in the single-object situation. However, this preference is not evident in the third condition (37° vs. 60°, Fig. 3 E).

To assess how flies select one or the other object, we analyzed when and where flies start fixating the object and what might have induced selection of one of the objects. We defined the start of object fixation as the onset of the time window where at least one of the object’s edges has been kept in the frontal visual field, i.e. within ± 30° relative to the midline of the animal, for at least 100 frames (^~^1.11s). In most cases, the lock-on is observed early after leaving the walkway (Fig. 3). The time between leaving the walkway and the fixation lock-on in presence of a single object, is very similar irrespective of object position, with only a slight tendency to take longer if the object is more lateral. When two objects are present the time to fixation onset is similar irrespective of object position and is only slightly larger compared to the single object experiments (Fig. 4 A). Fixation lock-on is located in many cases on the walkway or immediately upon leaving it irrespective of object position; only in few cases are fixation onsets located at some distance away from the walkway (Fig. 4 B).

To understand how blowflies select the object they will approach, we determined whether the animals tend to select the object they saw first when walking along the walkway. We calculated the phi-coefficient as a measure of the correlation between an object seen first and the object eventually approached. When the two objects are in a frontal position (37° and 60°) we find no correlation between seeing one of them first and approaching it (Phi-coefficient = 0.14). Furthermore, if one of the objects is placed at 90° it is never seen first but is still approached roughly in a third of the times (compare Fig. 3 A to Fig. 3 E - G). Hence, whether an object is seen first is unlikely to be a major determinant of object choice.

Does the mean walking speed of blowflies change after the animal started fixating the object? Across all experiments, the mean walking speed before and after fixation lock-on does not differ systematically (Fig. 5), suggesting that flies do not change their speed once they apparently decided to approach an object.

### Modelled walking behavior

In our experiments, walking blowflies seem to make remarkably quick choices, often starting to approach their goal early after leaving the walkway (Fig. 4 A). How can this behavior be explained? In tethered flight Poggio and Reichardt (1973) explained object choice as the result of each of the objects independently leading to a torque component. The torque caused by each object (‘object response’) follows a characteristic curve with the torque induced by an object depending on its azimuthal position in the visual field (Reichardt and Poggio 1976). The object responses to each individual object are assumed to be summed and added to spontaneous torque fluctuations. We analyzed whether this additive fixation model (AFM, Fig. 2) can explain the quick decisions we observed for freely walking blowflies.

To account for the spontaneous fluctuations in the walking direction (see Fig. 3), we added the object-induced response to angular velocity fluctuations based on the walking tracks of flies in our arena without any object (Fig. 3 A; see Methods and Monteagudo Ibaretta 2020). Since the walking speed of blowflies did not change much after fixation lock-on as well as with the different stimulus constellations, we set a constant modelled walking speed *v* = 6.4 cm/s, the average speed of blowflies recorded across all object constellations. With these parameter constellations we could fit the spontaneous walking behavior of flies quite well (compare Fig. 3 A and Fig. 6 A)

When confronted with two objects, one at 37° and one at 60°, the trajectories generated by the AFM consist mostly of rather smooth curves that initially head towards the midpoint between the objects and later bend towards one of them (Fig. 6 B). This behavior is to be expected, based on previous literature (Reichardt and Poggio 1976), as the tendency to turn towards one object is cancelled initially by the tendency to turn towards the other, i.e. when the retinal positions of the objects are relatively close to each other. However, this model performance does not match the behavior observed in walking blowflies in the same situation (Fig. 3 E), in which in most cases the animals started moving towards one of the objects shortly after leaving the walkway.

In order to generate the early decision to walk towards one of the objects after the animal leaves the walkway, we hypothesized that the animal may ignore for some time one of the objects, i.e. does not react to it, as has been described for flying fruit flies (Wolf and Heisenberg 1980, Sareen et al. 2011). Therefore, we elaborated the AFM into our stochastic attention model (SAM) by adding a random process deciding whether the object response is taken into account for generating the overall behavior in each update step of the model (see Methods).

The trajectories generated by the SAM (Fig. 6 C-H) describe mostly straight or slightly curved paths that often lead to the object in the one-object constellation, or to one of the objects when two are present. Most important, the SAM accounts for the early decision to approach an object in a similar way as observed in the experimental data. Moreover, a substantial number of trajectories do not lead to an object and even a few modelled trajectories seem to change direction after the simulated animal first moved for some time on a fixation course. In addition, the model consistently generates a preference for individual frontal and fronto-lateral objects, i.e. objects at 37° or 60°, over lateral objects, as observed in walking blowflies. This preference is also observed when the model animal can choose between two objects at 37° or 60° versus one object at 90°. However, the model also produces a preference for an object at 37° over one at 60° in both the one-object and the two-object constellation, a result differing quantitatively from its experimental counterpart (Fig. 3).

Note that the data shown in Fig. 6 has the same number of trajectories per condition as the experimental data (100 walks). By re-running the simulation with different random seeds, a smoother estimate of the distributions shown in the polar histograms can be achieved (cf. Fig. S2).

The time to fixation lock-on (Fig. 7 A) is fairly quick in the one-object constellation if the object is at 37° and increases slightly for more lateral positions of the object. In the two-object constellation the effect is the same, with the time to fixation onset being shorter when the objects are at 37° and 60° and somewhat longer for object positions at 37° and 90° and longest for a combination of 60° and 90°. Although this position dependence is not very strong, it differs to some extent from the experimental data. Also, the time to fixation onset tended to be shorter in the experiments even when compared to the shortest times observed in the model simulations.

**Figure 7:**
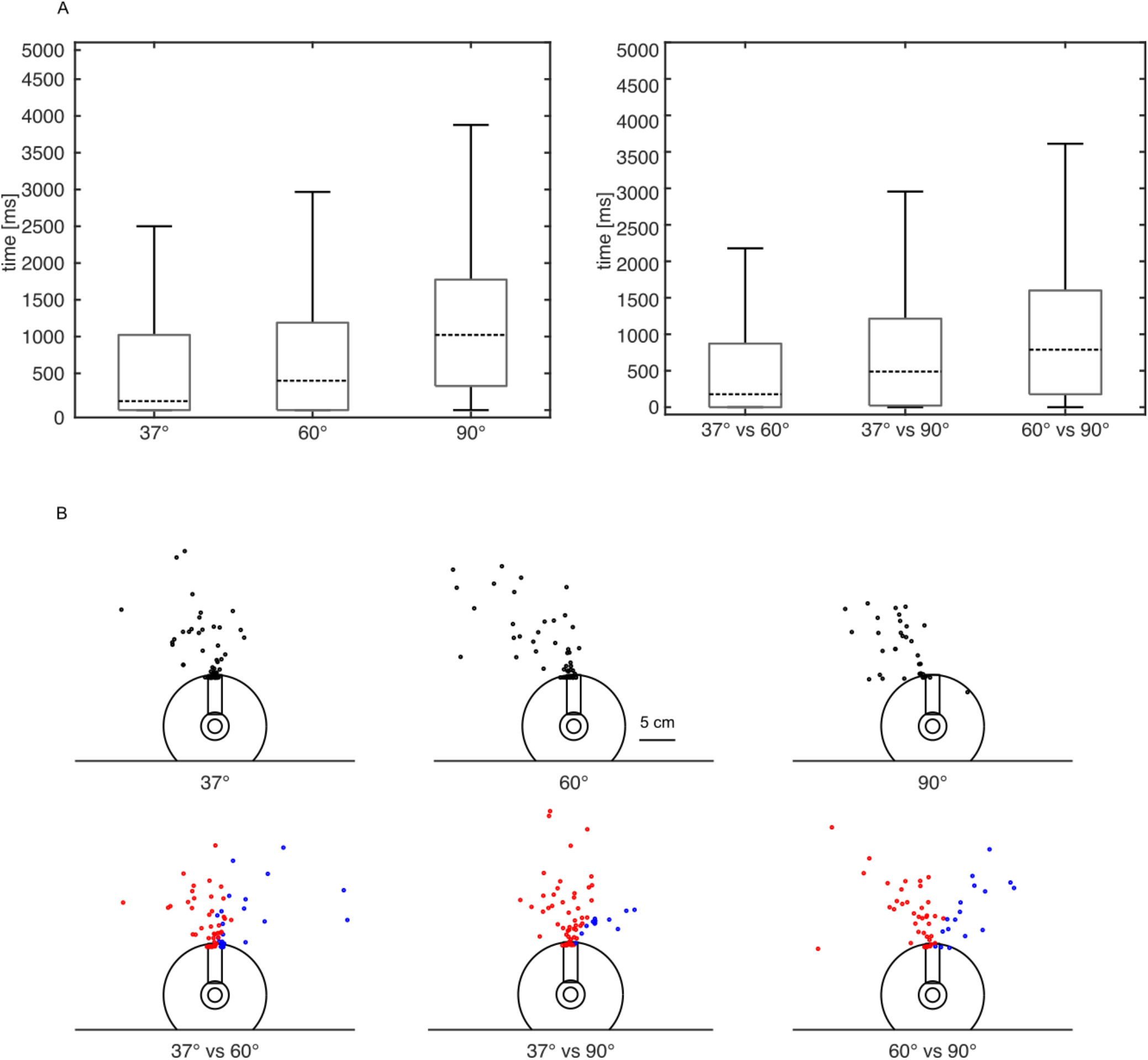
Analysis of object lock-on of the stochastic attention model (SAM). A) Boxplots of the time between leaving the walkway and object lock-on and B) Location of lock-on moment. Upper row: single objects. Lower row: two objects, red dots indicate walks approaching the object more frontal to the walkway, blue dots the one more lateral.

We also plotted the locations of the fixation onset for the modelled blowflies (Fig. 7 B). For both the one- and two-object constellation, fixation onset is located close to the start of the trajectory at the end of the walkway. However, as a consequence of the wider spread of the time to fixation in the model data compared with the experimental counterpart, the locations of fixation onset are spatially more spread, especially when one object is at 90° (compare Fig. 7 B and Fig. 4 B). Irrespective of these quantitative differences, the SAM matches the experimentally determined behavior in our two-object paradigm quite well.

## Discussion

The research literature on object-related orientation behavior of flies reports two seemingly contradictory findings. On the one hand, a reflex-like fixation mechanism was proposed that continuously adds the responses to individual objects (Reichardt 1973, Reichardt and Poggio 1976). On the other hand, open-loop experiments with tethered flying flies suggest the presence of selective attention and active choice processes under symmetrical stimulation with moving objects (Wolf and Heisenberg 1980).

These conclusions were mainly based on experiments on tethered flying flies in situations where translation velocity could not be determined and, thus, did not affect the visual input, implying that the objects were virtually placed at infinity. Therefore, we addressed the issue of object-related orientation with a choice paradigm in which blowflies could walk freely towards an object. When confronted with two objects, free-walking flies show behaviors which might be interpreted as active decisions to approach either object. The flies seemed to quickly select the target approached. The initial azimuthal positions of the potential targets have an influence on their attractiveness but have little impact on the time needed to select the object to approach. However, which of the objects was seen first does not play an obvious role in target selection.

To investigate possible mechanisms underlying this selection process, we developed a computational model qualitatively reproducing the animals’ performance. This model revealed that we cannot explain the apparent quick decision process and therefore cannot account for our behavioral data without including an attention-like mechanism. However, the stochastic attention mechanism we implemented does not explicitly assume a decision process or memory.

By observing behavior, one cannot pinpoint exactly when an animal draws a decision, but rather has to rely on visible behavioral indicators, such as the onset of a turn, as a proxy for decision. With the spatial layout of the setup, forcing the animals to walk over a narrow walkway into the setup of the possible fixation targets, we tried to harmonize the initial walking directions and retinal object positions. For analysis purposes we defined the time at which the animal leaves the walkway as the reference point in time. At this point all objects were visible for the animal in all conditions. We conclude from our observations that walking blowflies show a preference for objects depending on their initial azimuthal position, with frontal objects being preferred over lateral ones, both when only a single object is presented and when the flies are allowed to choose between two. Despite reacting to objects in an obvious manner, blowflies did not walk towards any object in a sizable proportion of walks for all tested object configurations and walked in other directions.

Once flies start fixating their goal, they tend to stick to this apparent decision. The short time interval between leaving the walkway and lock-on in all tested object configurations suggests that flies take a very similar amount of time to respond to the objects regardless of their position and that, even in a two-alternative choice situation, they only take slightly longer to decide. Thus, despite inevitable limitations in our methodology, we can be confident that the retinal position of the objects has little effect on the time the animal takes to decide to approach one of them. This is in accordance with a previous study (Mronz 2004) reporting largely constant reaction times of walking *Drosophila* towards objects presented at different azimuthal positions in the frontal visual hemisphere (0°-90°), though the reaction times observed in this study were with around 1s much larger than the ones we observed in *Lucilia*. This difference could result from differences in the experimental paradigms or species used, as Mronz (2004) measured the reaction time of *Drosophila* to sudden position changes of bars, while in our paradigm the objects became visible to the blowfly slightly before the reference time point.

The time until fixation increases a bit compared to the single-object condition, when the fly is confronted with two objects. A delay when choosing between multiple objects has been described for flies in the context of attention (van Swinderen 2011), arguing that the presence of additional objects, referred to as distractors, draws attention away from the target objects. For *Drosophila,* in particular, it has been shown that the animal responds to only one of two stripe patterns moving in opposing directions in each half of the visual field, but that the response onset is delayed compared to the known response to a single rotating panorama (Tang and Juusola 2010).

We tried to find a parsimonious behavioral mechanism that could qualitatively reproduce the observed quick apparent object choice. It has been proposed that choice behavior between different objects is the result of the animal adding the turning responses generated by a static fixation characteristic to the available objects and spontaneous noise-like turning tendencies (Reichardt and Poggio 1976). We first implemented this mechanism in our additive fixation model (AFM). In simulations we found that this model fails to generate a quick fixation decision between objects being close to each other. The AFM can generate a choice between two objects only if these are sufficiently separated. Otherwise, both objects generate similar turning responses in opposite directions causing the simulated animal to walk towards an intermediate position, as has been described before during tethered flight (Reichardt and Poggio 1975). We conclude that, while we can find individual simulation runs of traces leading to an object from the start, in most cases the AFM is unable to generate a quick decision as we observed in walking blowflies.

Thus, we hypothesized that to quickly fixate an object in a two-object paradigm, it might be necessary for the animal to ignore one or the other object at least occasionally. This ability had already been concluded for *Drosophila* in a scenario involving two vertical objects in an open-loop paradigm (Wolf and Heisenberg 1980) and thus seems to be a reasonable assumption. The implementation in our stochastic attention model (SAM) assigns a probability to react to each object which depends on its azimuthal position. This extension of the model leads to qualitative reproduction of the quick responses observed in walking blowflies and thus revealed that ignoring part of the visual input for some short time window is sufficient to explain the observed behavior. This finding may immediately evoke the notion of attention, i.e. the ability to focus on parts of the visual input while ignoring the rest. Indeed, our two-object paradigm leads to conclusions reminiscent to the ones of studies used to showcase and study competitive attention (van Swinderen 2011, Nityananda 2016). In these studies, an animal had to respond to a visual stimulus while suppressing the response to the other.

For our SAM we simulated that each object had an independent probability to be taken into account or, conversely, to be ignored at each time step. The probability of attendance varied depending on the object’s azimuthal position, reaching a maximum for a frontal object. Our model does not need to keep track of internal states or memorize the decisions taken, the generated object fixation and decision-like processes are emergent properties. The SAM can reproduce features of the observed animal’s behavior: despite producing a robust fixation, it allows for significant proportions of modelled flies to not approach any of the objects as well as to occasionally switch to a different goal after apparently starting to walk towards an object, as observed in walking blowflies.

For two objects the choice mechanism implemented in the SAM effectively selects one of four response types in each time step of the simulation:

1. Stabilizing the fixation of an object when the more frontal object is the only one contributing to the yaw torque,
2. reorientation towards the other object if only the response to the more lateral object is ‘on’,
3. turning towards a compromise path when both objects contribute, or
4. ignoring both objects altogether.

By modulating the probability of the stochastic process depending on object position in a bell-shaped attention characteristic, the first state, stabilizing fixating the most frontal object, is selected with the highest probability, while the system still can reorient by randomly selecting one of the other states.

Multiple other factors affect choice and fixation behavior of flies apart from the azimuthal position of objects in the visual field. For example, *Drosophila* shows different preferences for bars depending on how wide they are (Wehner 1972), and *Lucilia* has different preferences for different colors (Fukushi 1989). The SAM could be extended to address other preferences by tuning the object response curve to objects of variable characteristics or by adapting the attention curve to reflect preferential attention based on other stimulus parameters than the azimuthal position.

## Acknowledgments

We thank Roland Kern for critically reading and helpful comments on the manuscript and Tim Siesenop for animal breeding.

## Supplementary Figures & Legends

**Figure S1:**
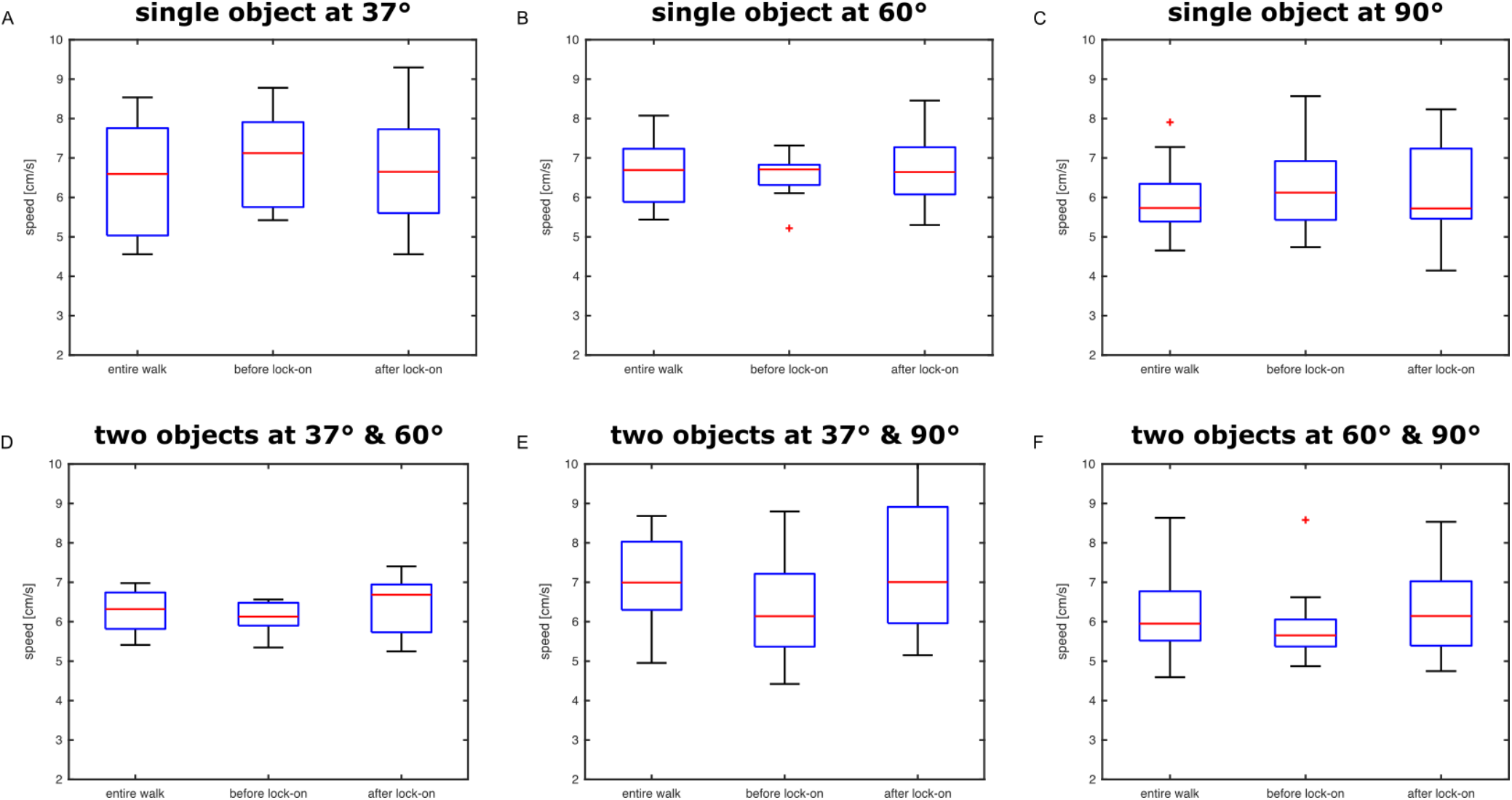
Walking speed of flies in different phases of the walk. Boxplots show median walking speed (red line), 25% and 75% quantile (box) and data range (whiskers, where “outliers” are indicated as ‘+’). Walking speed distributions are visualized for the entire walk and the segments before and after lock-on for each object condition. Upper row (A-C) shows single-object conditions, Lower row (D-F) conditions with two objects.

**Figure S2:**
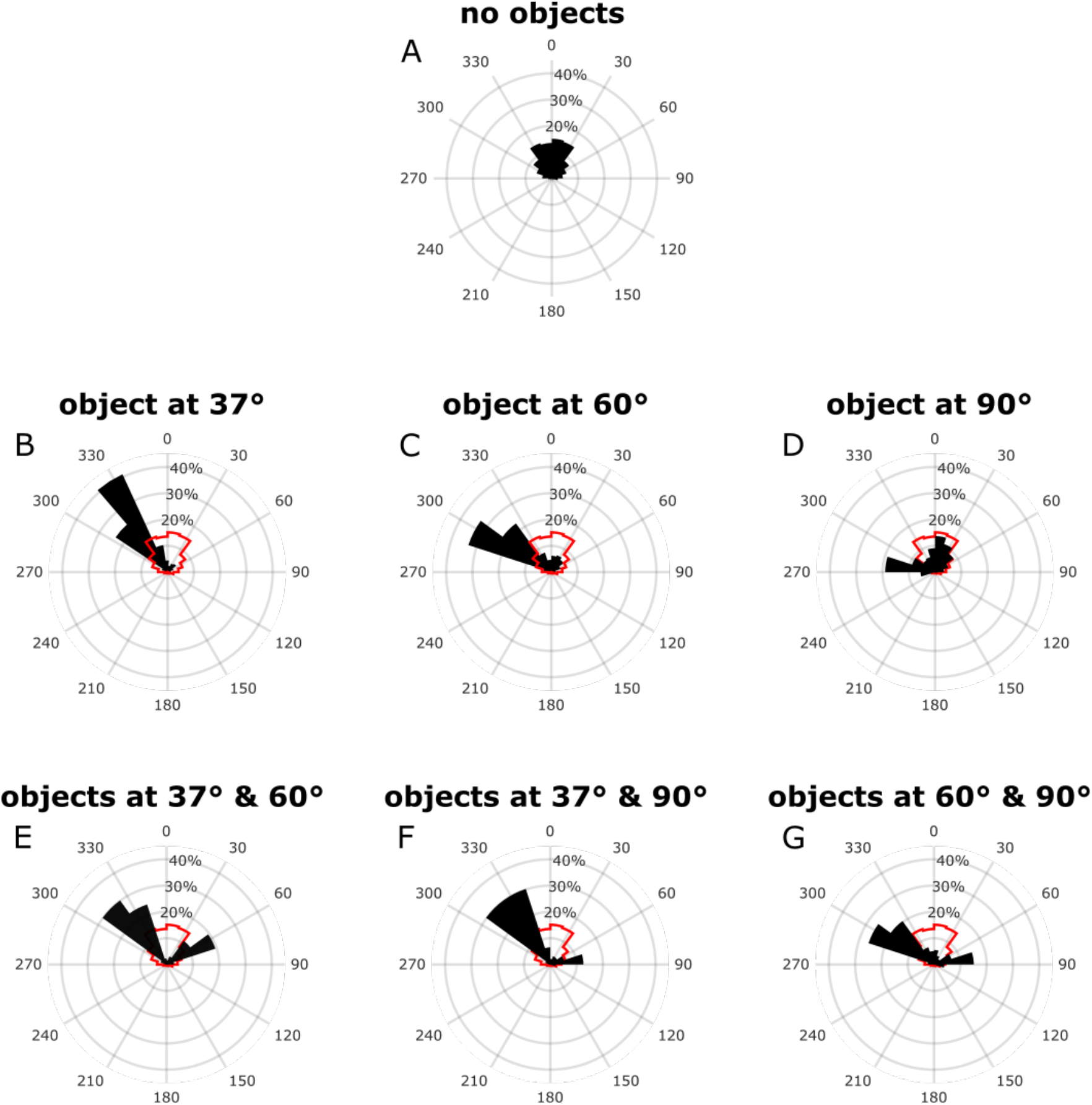
Walking trajectories generated by the model. Data for 1000 walks per condition: Simulations as in Fig. 6 repeated with 10 different random number seeds. A) Data resulting from spontaneous turns in absence of any object. B-D) Data generated by the SAM for single objects. E-G) Data generated by the SAM when tested with two objects. The polar histograms show the distribution of positions observed when the model trajectories crossed a registration circle with 20 cm radius around the end of the walkway. Red silhouettes in B - G repeat the distribution of A for comparison.

## References

Fukushi, T. (1989). Learning and discrimination of coloured papers in the walking blowfly, Lucilia cuprina. J. Comp. Physiol. A 166, 57–64.

Horn, E., & Wehner, R. (1975). The mechanism of visual pattern fixation in the walking fly, Drosophila melanogaster. J. Comp. Physiol. 101, 39–56.

Horn, E. (1978). The mechanism of object fixation and its relation to spontaneous pattern preferences in Drosophila melanogaster. Biol. Cybern. 31, 145–158.

Monteagudo Ibaretta, J (2020). Object directed behavior of walking blowflies. PhD thesis, Bielefeld, Germany: Universität Bielefeld. DOI:10.4119/unibi/2943426

Mronz, M. (2004). Die visuell motivierte Objektwahl laufender Taufliegen (Drosophila melanogaster) - Verhaltensphysiologie, Modellbildung und Implementierung in einem Roboter. PhD thesis, Würzburg, Germany: Universität Würzburg, urn:nbn:de:bvb:20-opus-11748

Nityananda, V. (2016). Attention-like processes in insects. P. Roy. Soc. B: Biol. Sci. 283, 20161986.

Poggio, T., & Reichardt, W. (1973). A theory of the pattern induced flight orientation of the fly Musca domestica. Kybernetik 12, 185–203

Reichardt, W. (1973). Musterinduzierte Flugorientierung. Naturwissenschaften 60, 122–138.

Reichardt, W., & Poggio, T. (1975). A theory of the pattern induced flight orientation of the fly Musca domestica II. Biol. Cybern. 18, 69–80.

Reichardt, W., & Poggio, T. (1976). Visual control of orientation behaviour in the fly: Part I. A quantitative analysis. Q. Rev. Biophys. 9, 311–375.

Sareen, P., Wolf, R., & Heisenberg, M. (2011). Attracting the attention of a fly. PNAS 108, 7230–7235.

Schuster, S., Strauss, R., & Götz, K. G. (2002). Virtual-reality techniques resolve the visual cues used by fruit flies to evaluate object distances. Curr Biol 12, 1591–1594.

Tang, S., & Juusola, M. (2010). Intrinsic activity in the fly brain gates visual information during behavioral choices. PLoS One 5, e14455.

van Swinderen, B. (2011). Attention in drosophila. Int. Rev. Neurobiol. 99, 51–85.

Warzecha, A.-K. & Egelhaaf, M. (2000). Response latency of a motion-sensitive neuron in the fly visual system: dependence on stimulus parameters and physiological conditions. Vis. Res. 40, 2973–2983.

Wehner, R. (1972). Spontaneous pattern preferences of Drosophila melanogaster to black areas in various parts of the visual field. J. Insect Physiol. 18, 1531–1543.

Wolf, R., & Heisenberg, M. (1980). On the fine structure of yaw torque in visual flight orientation of Drosophila melanogaster. J. Comp. Physiol. 140, 69–80.

